# High throughput measurement of *Arabidopsis thaliana* fitness traits using deep learning

**DOI:** 10.1101/2021.07.01.450758

**Authors:** Peipei Wang, Fanrui Meng, Paityn Donaldson, Sarah Horan, Nicholas L. Panchy, Elyse Vischulis, Eamon Winship, Jeffrey K. Conner, Patrick J. Krysan, Shin-Han Shiu, Melissa D. Lehti-Shiu

**Author notes:** Authors for correspondence: Shin-Han Shiu, Lab website: https://shiulab.github.io, Melissa D. Lehti-Shiu. Joint first authors (Peipei Wang, Fanrui Meng).

## Abstract

- Revealing the contributions of genes to plant phenotype is frequently challenging because the effects of loss of gene function may be subtle or be masked by genetic redundancy. Such effects can potentially be detected by measuring plant fitness, which reflects the cumulative effects of genetic changes over the lifetime of a plant. However, fitness is challenging to measure accurately, particularly in species with high fecundity and relatively small propagule sizes such as *Arabidopsis thaliana*.
- An image segmentation-based (ImageJ) and a Faster Region Based Convolutional Neural Network (R-CNN) approach were used for measuring two Arabidopsis fitness traits: seed and fruit counts.
- Although straightforward to use, ImageJ was error-prone (correlation between true and predicted seed counts, r^2^=0.849) because seeds touching each other were undercounted. In contrast, Faster R-CNN yielded near perfect seed counts (r^2^=0.9996) and highly accurate fruit counts (r^2^=0.980).
- By examining seed counts, we were able to reveal fitness effects for genes that were previously reported to have no or condition-specific loss-of-function phenotypes.
- Our study provides models to facilitate the investigation of Arabidopsis fitness traits and demonstrates the importance of examining fitness traits in the study of gene functions.

## Introduction

A major goal of biology is to understand the molecular basis for the development of organisms and their adaptation to different environments (Mcdonald, 1983). One approach is to evaluate the effects of genetic variants on phenotypes. However, it is often challenging to investigate such effects because gene functions may be masked by genetic redundancy (Bouché & Bouchez, 2001; Sun et al., 2012) and/or be condition specific (Hirsch *et al*., 1998; Meissner *et al*., 1999). Moreover, the physiological and/or developmental changes caused by loss of gene function may be too subtle to detect. This challenge can be alleviated by measuring the effects of genetic variations on fitness (i.e., the ability of an individual to survive and reproduce) because it reflects the cumulative effects of genetic changes over the lifetime of a plant. Accurate estimates of fitness are therefore valuable for several fields of study, including plant genetics, evolution, and plant breeding.

Among fitness measures, the most direct measure is the number of progeny produced (Thomson & Hadfield, 2017). In *Arabidopsis thaliana*, a predominantly selfing plant, the total number of seeds produced per plant is a particularly good estimate of fitness because it incorporates both male and female contributions. However, because Arabidopsis seeds are small (areas ranging from ∼0.1–0.2 mm^2^; Jahnke *et al*., 2016) and produced in large numbers (up to thousands of seeds per plant; Boyes *et al*., 2001; Morales *et al*., 2020), it is difficult to obtain accurate seed counts. As a consequence, fruit (silique) number (Busoms *et al*., 2015) and total fruit length (Roux *et al*., 2004; Busoms *et al*., 2015; Kerwin *et al*., 2015) are often used to measure fitness. Both measures are correlated with seed production, but fruit number is not perfectly correlated with seed number (e.g., r^2^=0.960, Mauricio & Rausher, 1997) and correlations in fruit length are highly variable across studies, ranging from and r^2^=0.988 (Roux *et al*., 2004) to r^2^=0.256 (Gnan *et al*., 2014). In addition, fruit numbers (up to 450 per plant; Hamidinekoo *et al*., 2020) are typically counted manually, and these counts can be error prone. Thus, to better measure fitness, both fruit and seed numbers should be evaluated using methods that are not hindered by propagule size or number.

Several programs have been designed to increase the efficiency and accuracy of seed analyses. Some are aimed at measuring the properties of individual seeds (e.g., size and shape) rather than obtaining high throughput seed counts (Herridge *et al*., 2011; Tanabata *et al*., 2012; Moore *et al*., 2013). These approaches typically require that seeds be separated before imaging, which increases the time needed for processing. Other systems have been designed to separate seeds mechanically. For example, the *pheno*Seeder device designed by Jahnke *et al*. (2016) separates seed using a “pick-and-place” robot and using a large-particle flow cytometer to separate seeds Morales *et al*. (2020) were able to count an average of 12,000 seeds per hour with high accuracy (relative error < 2%). A similar number of seeds, approximately 10,000, can be processed per hour using the BELT imaging system combined with the phenoSEED algorithm, which acquires images of individual seeds as they pass through an imaging chamber (Halcro *et al*., 2020). Nonetheless, a drawback of these methods is that they require specialized equipment, hindering their widespread adoption.

Another approach that has been increasingly used in plant biology for applications such as measurement of fitness traits is machine vision, the application of machine learning algorithms to image analysis (Mochida *et al*., 2019). Deep learning approaches, in particular Convolutional Neural Network (CNN)-based frameworks, have been developed to detect vastly different objects (from cars to plant seeds) in images. For example, the seeds of rice, lettuce, oat, and wheat, were detected with 96% recall and 95% precision using Mask R-CNN (Toda *et al*., 2020). However, the detection of much smaller objects using CNN-based approaches remains challenging (Cao *et al*., 2019). This is likely because CNNs create low-level abstractions of the images, and if the objects are too small, the resulting abstractions are too simple to be used to distinguish whether the object is present or not. Although the CNN-based models developed by Toda *et al*. (2020) detected seeds with high accuracy, the smallest seeds tested were lettuce seeds, which have areas ranging from 1.5–3.6 mm^2^ (Penaloza *et al*., 2005) and are ∼10 times larger than Arabidopsis seeds. Another consideration is that the most convenient way to count all the seeds from an Arabidopsis plant, which can produce thousands of seeds (Boyes *et al*., 2001; Morales *et al*., 2020), would be to put all the seeds in a single image, thus resulting in a relatively small ratio of seed size to image size. However, due to the small image size (1024 × 1024 px^2^ or 2000 × 2000 px^2^) used in Toda et al. (2020), the ratio of seed size to image size was relatively large (>5000 px^2^ per barley seed), and the number of seeds that could be included in an image was also limited. Therefore, it is important to assess how well the CNN-based approaches perform in detecting objects as small as Arabidopsis seeds in an image containing thousands of them.

CNN-based approaches have also been used in fruit counting. For example, using R-CNN, wheat spikes can be detected, counted, and analyzed to estimate yield based on images captured in the field (correlation between true and predicted counts: r^2^=0.93 with slope of 1.01; Hasan *et al*., 2018). Starting from two pre-trained models (ResNet and ResNext), Afonso *et al*. (2020) applied the Mask R-CNN approach to detect and count tomato fruits from images captured in the greenhouse, obtaining an F1 of 0.94 when fruits were only partially overlapping with each other. DeepPod, developed using the LeNet and DenseNet-Basic models as a starting point, effectively counts Arabidopsis fruits but results in a high number of false negatives when there are many fruits (r^2^=0.90 with a slope of ∼0.70; Hamidinekoo *et al*., 2020). In addition, the inflorescences need to be harvested when the fruits are still green, preventing the harvesting of seeds for future propagation or analysis. Thus, it is important to develop tools or models to detect and count mature fruits when seeds need to be saved for future experiments. Because Arabidopsis fruits shatter easily when dry, such tools should ideally be able to count fruits at different stages, including intact fruits and those that have already dehisced and released seed, when it is necessary to score plants grown to maturity (Conner & Rush, 1997).

In this study, we evaluated two existing approaches—segmentation with ImageJ (Schneider *et al*., 2012) and a deep learning approach, Faster R-CNN (Ren *et al*., 2017)— for counting all the seeds an Arabidopsis plant produces in a single image. We also applied Faster R-CNN to count fruits in whole plant images captured after seeds were mature. To facilitate seed and fruit counting in diverse images, we established models using input images with varying resolution, contrast, brightness, and blurriness. The final seed and fruit models are provided and can be readily used by the research community. Finally, we used our pipeline to count seeds for loss-of-function mutants of three genes that were previously reported to have no obvious phenotype, a condition-specific phenotype, or no reported phenotype. We showed that mutation of these genes does indeed affect fitness, illustrating the importance of measuring fitness traits and the utility of our pipeline in the investigation of gene functions.

## Materials and Methods

### Plant materials

T-DNA insertion mutants in the *Arabidopsis* Col-0 background and wild-type (WT) Col-0 controls were used for training seed and fruit counting models. Information about these lines is provided in **Tables S1 and S2**. Fitness data are reported for T-DNA insertion mutants of *PURPLE ACID PHOSPHATASE 2* (*AtPAP2*, *At1g13900*, SALK_013567C), *KINESIN 7.4* (*AtKIN7.4*, *At4g39050*, SALK130788C), and *RECEPTOR DEAD KINASE 1* (*AtRDK1*, *At4g34220*, SALK_112336). For these mutants, multiple homozygous mutant and WT sibling plants were identified by PCR with gene-specific primers (two to seven plants per genotype; **Table S3**). Seeds harvested from these independent lines (referred to as sublines) were planted (*n* = 5–20 per subline, total *n* ≥ 40 per genotype) for comparison of fitness between mutants and WT, and each mutant was compared with its WT sibling. This was done to reduce the chance that observed fitness effects were due to other undetected T-DNA insertions.

For fitness analysis and seed counting, seeds were sown in 200-cell flats filled with Arabidopsis mix (1:1:1 SureMix, vermiculite, and perlite), stratified for 5–7 days in the dark at 4°C, then transferred to a growth chamber and grown under a 16-h light/8-h dark cycle with a light intensity of 110–130 μmoles/m /s at 21°C. Genotypes were randomly assigned to cells within a flat. Seedlings were thinned to one per cell after 1 week. Plants were watered two to three times per week and grown to maturity. Intact fruits were transferred to glassine envelopes and allowed to dry for at least one month before counting seeds.

For fruit counts, seeds were sown in 200-cell flats filled with MetroMix 360, stratified for 7 days in the dark at 4°C, then grown under a light intensity of ∼150 μmoles/m /s at 21°C. Genotypes were randomly assigned to cells within a flat. Seedlings were thinned to one per cell after 1 week. After five weeks (23 November 2018) flats were transferred to a plot at Kellogg Biological Station. Plants were removed from the field on 23 May 2019. Plants were cut at the base and imaged as described below.

### Seed image scanning

Seeds were scattered on the lid of a 60-mm petri plate, which was then tapped with fingers to ensure that seeds did not cover each other. For some plate lids counted with ImageJ, seeds were separated from each other and moved away from the edges of the lid using forceps. The lids were placed in a template made from white acrylic (295 mm × 210 mm × 10 mm, **Fig. 1a**) on a standard desktop scanner. This template was designed to be the same height as the plate lid to minimize artifacts resulting from light loss. Twelve 60-mm diameter holes (four rows × three columns) were cut into the template using a laser cutter. There was 6 mm between holes in each row and 10 mm in each column. The template was aligned to the top right corner of the scanner before every scan because even small changes (<1 cm) in the location of the template required changing the parameters for the circular search regions in the ImageJ algorithm (described in the following section). Seeds were scanned at 1200 dots per inch and in 24-bit color, and scans were saved as jpeg files.

**Fig. 1.**
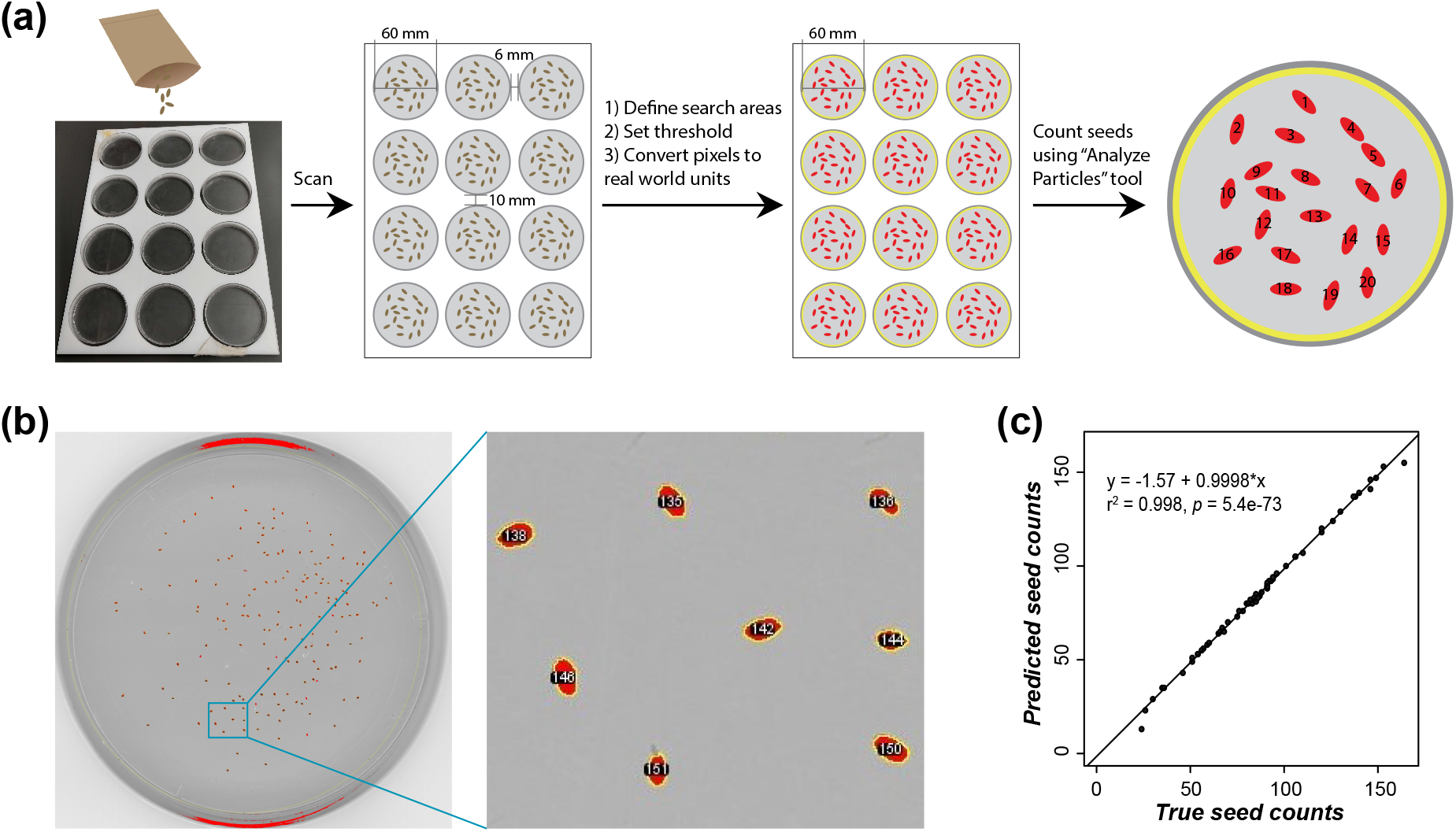
Workflow and performance for seed counting using ImageJ when seeds were deliberately separated. (**a**) Workflow. Seeds from 12 different plants were scattered and manually separated from each other on the lids of 12 petri plate lids, which were placed in a template and scanned. Twelve search areas, each with a diameter of 60 mm (yellow circles), were predefined. A threshold was applied by selecting pixels with intensities between 50 and 140 to separate the seed areas (red) from the background. Then pixels were converted to real-world distance units in mm. The “Analyze Particles” tool was used to detect and count the seeds. (**b**) An example of an image with detected seeds (left) and an enlarged image showing the seeds (right). Red region with number: individual detected seed area. (**c**) Correlation between true and predicted seed counts using ImageJ when seeds were deliberately separated.

### Seed image processing and counting with ImageJ

The ImageJ (version 1.52a, https://imagej.nih.gov) workflow is shown in **Fig. 1**. Scanned images were first converted into 8-bit grayscale bitmap format with the im.convert(‘L’) method in the Python Imaging Library (https://pythonware.com/products/pil). An ImageJ macro was written to automate seed counting. First, 12 circular search regions per scanned image were defined to confine the search to the 12 plate lids containing the seeds. Second, a threshold was applied by selecting pixels with intensities between 50 and 140, and pixels were converted to real world units using a predefined line covering a known distance (the distance across the plate lid, 60 mm). Third, seeds within the circular search regions were counted using the “Analyze Particles” tool. Only above-threshold regions larger than 0.06 mm^2^ and with a circularity value (calculated as C=4π × [area/perimeter]) of 0.25 to 1 were counted. The image conversion program and the ImageJ macro were combined into a Windows batch script (available in our Github repository, see **Data availability**), in which a for-loop was used to quickly count seeds for images in sequence. It took approximately 5 min to fully process 10 images.

### Seed image processing and counting with Faster R-CNN

Each scanned image was first split into 12 sub images; each sub image contains a single plate lid and is referred to as a “whole-plate image”. In the initial Faster R-CNN modeling trial, each whole-plate image was split into four quarter-plate images. Seeds on the quarter-plate images were manually annotated using LabelImg v1.8.1 (Tzutalin, 2015). The ground truth annotations, i.e., the coordinates of the bottom-left (x_min_, y_min_) and upper-right (x_max_, y_max_) corners of each annotated seed area, were saved in extensible markup language (xml) format, and then converted to comma-separated values (csv) format. For images used for model training, the csv files were further converted to the TFrecord format.

Faster R-CNN, performed using Tensorflow object detection API (Huang *et al*., 2017) and implemented in Tensorflow v1.13.2 (Abadi *et al*., 2016) in python v3.6.4, was used for seed detection. Faster R-CNN combines the generation of region proposals (i.e., circumscribing the areas of interest, a regression problem) and their classification (i.e., in our case, the object is a seed or not) into a single pipeline (Ren *et al*., 2017). In Faster R-CNN, images were first processed by a feature extractor (Inception V2; Szegedy *et al*., 2016), and the resulting feature maps were used to predict bounding boxes (referred to as proposals) containing images of individual seeds (left panel in **Fig. S1**); then these proposals were used to crop features from the feature maps (right panel in **Fig. S1**). These cropped features were subsequently used for classification and bounding box regression.

To speed up the training process, a pre-trained model (<u>faster_rcnn_inception_v2_coco</u>) was used as a starting point. To optimize Arabidopsis seed detection, three hyperparameters were tested: proposals—the number of detected regions (i.e., seeds) in an image; scales—relative sizes of detected regions; and aspect ratios—shapes of the detected regions (**Fig. 2a** and **Fig. S1**). For the hyperparameter space tested see **Table S4.** Each model was run on three graphics processing units (GPUs) for 40,000 steps using the Adam optimizer, with a batch size of five (five random images were used to train the model for each step) and a learning rate of 0.0002. A model was saved every 10 minutes during the run as a model checkpoint.

**Fig. 2.**
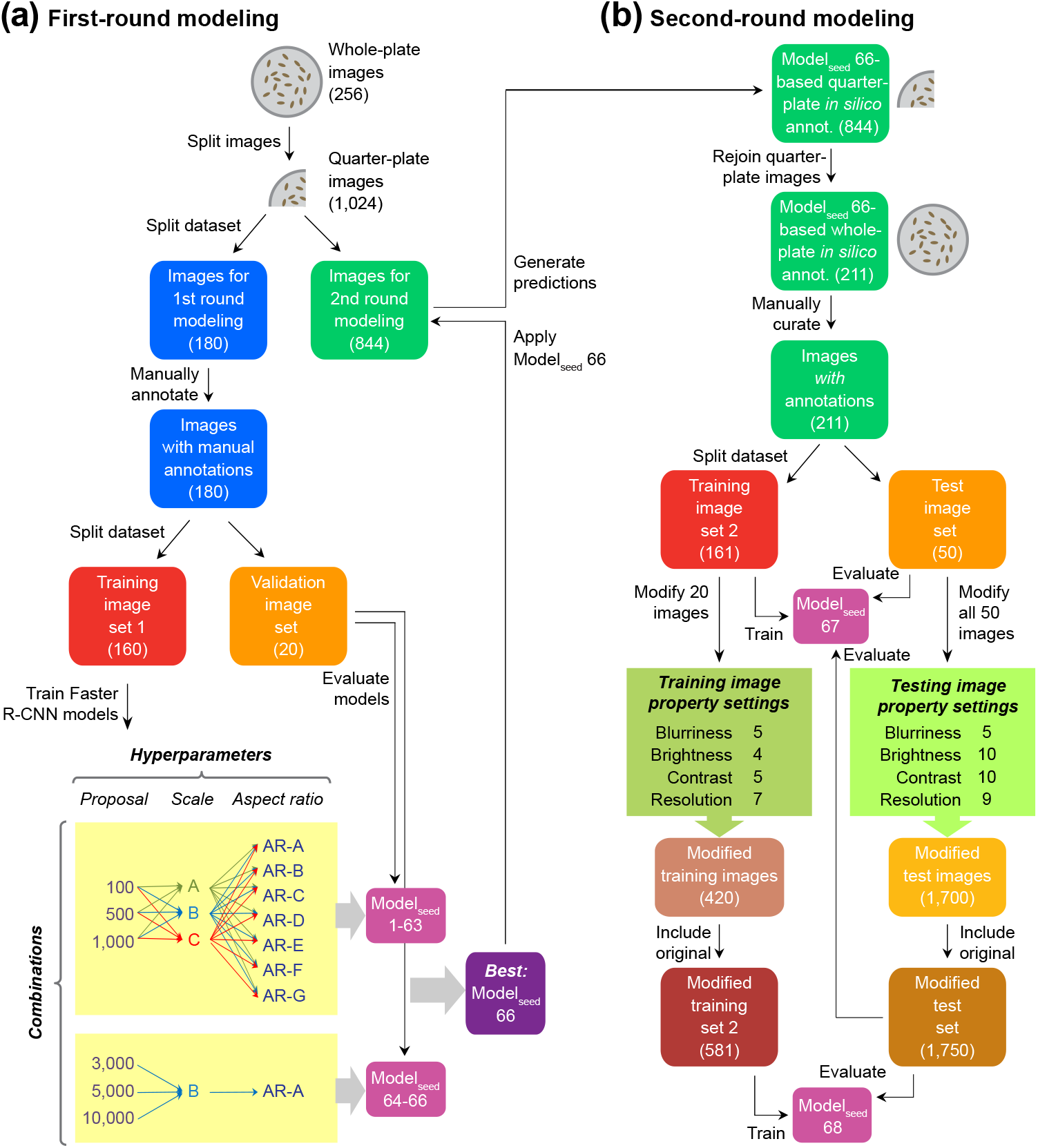
Workflow for building Faster R-CNN-based seed counting models. (**a**) First-round modeling for enriching annotated seed labels. Each of the 256 whole-plate images was split into four quarter-plate images. Among the 1024 quarter-plate images, 180 were used in first-round modeling, and the remainder (844) were used in second-round modeling described in (**b**). Seeds in the 180 quarter-plate images were manually annotated, and then these annotated images were further split into training set 1 (160) and a validation set (20) to train and evaluate models, respectively. Sixty-three combinations of three hyperparameters (i.e., 3 proposal numbers × 3 scales [A, B, and C] × 7 aspect ratios [AR-A through G]; for scale and aspect ratio values see **Table S4**) were used to build 63 models. The optimal scale (B) and aspect ratio (AR-A) were selected based on the model performance on validation set images (**Fig. S2**). An additional three models (Model_seed_ 64–66) were built using scale B, AR-A, and three larger proposal values, and the final best model, Model_seed_ 66, with 10,000 proposals, was applied to the 844 quarter-plate images reserved for second-round modeling to generate *in silico* seed annotations. (**b**) Second-round modeling. The 844 quarter-plate images with seed predictions from Model_seed_ 66 were rejoined together to reconstruct 211 whole-plate images with *in silico* seed annotations, which were then manually curated and used as ground truth seed annotations. Model_seed_ 67 was built using 161 (training set 2) out of the 211 annotated images with the same hyperparameters used in Model_seed_ 66, and was evaluated using the test set (50 independent images not used for modeling) and the modified test set (i.e., the 50 independent test set images plus 1,700 images modified from the test set images that had different image properties [blurriness, brightness, contrast, and resolution values]). For data augmentation, the image properties of 20 images from training set 2 were modified, and the resulting 420 images were combined with training set 2 (161 images), resulting in 581 images (modified training set 2), which were used to build Model_seed_ 68. The modified test set was used to evaluate the performance of Model_seed_ 68.

To evaluate the model performance for hyperparameter tuning, validation set images—seeds in these images were manually annotated but not used in the model training—were fed into the frozen models in jpeg format, and model predictions were output in csv files containing the coordinates of the bottom-left and upper-right corners of each predicted seed area. Coordinates of ground truth annotations in validation set images were compared with those of predicted seed areas using the measure IoU, which is defined as the intersection (I) over (o) the union (U) of a ground truth area and a prediction area. A seed was regarded as correctly detected if its IoU was ≥ 0.5. An IoU < 0.5 was considered to be a misprediction. The F-measure (F1) score was calculated as a measure of performance for a model as follows: 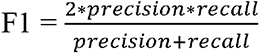, where precision 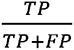 and recall 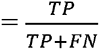. TP (true positive) is the number of correctly detected seeds, and FP (false positive) is the sum of the number of predicted seed areas that contain no seeds and the number of predicted seed areas minus 1 (if the ground truth area was detected more than once). FN (false negative) is the number of seeds with a maximum IoU < 0.5 for any predicted area (i.e., the ground truth seed area was not detected).

For the hyperparameter tuning, we found that models with 100 proposals (Model_seed_ 1–21) had the least accuracy, with F1 < 0.750 (**Fig. S2a**), because there were more than 100 seeds in most of the validation images (**Table S5**). In contrast, models with 500 (Model_seed_ 22–42) and 1000 proposals (Model_seed_ 43–63) had F1 values around 0.970 (**Fig. S2b,c**). For models with 500 or 1000 proposals, scale-B and scale-C had higher F1 values than scale-A, but there were no differences between the F1 values of the seven different aspect ratios (**Fig. S2b,c**). The combination of scale-B and aspect ratio-A was used for downstream model building because the computational efficiency of this combination was the highest among all combinations (**Fig. S3**). Three additional models were established based on scale-B and aspect ratio-A with 3000, 5000, and 10,000 proposals (Model_seed_ 64–66), but F1 values did not improve over those of Model_seed_ 22– 63 (**Fig. S2d**). Even though building Faster R-CNN models with a higher number of proposals requires more computational resources, we opted to use 10,000 proposals (Model_seed_ 66) for further model building to allow detection of a large number of seeds in an image.

### Fruit image capturing and counting with Faster R-CNN

Each dry Arabidopsis plant was placed on a pink paper background and photographed with an iPhone 8 smartphone. The images were saved in jpeg format with dimensions of 3024 × 4032 pixels. Fruits in the images were manually annotated, and the annotated coordinates were then converted to the csv and TFrecord formats, as conducted for the seed images. The same pre-trained Faster R-CNN model used for seed counting was used to build the fruit counting models, and the same three hyperparameters were tuned to optimize the model performance but with a different hyperparameter space (**Table S6**). For each hyperparameter combination, a model was saved after 6000 steps, when the performance had converged. A final model was established using hyperparameters selected based on performance on the validation set images.

## Results

### Seed counting using ImageJ

Because the “Analyze Particles” function of ImageJ is widely used for analysis of seed morphology (Cervantes et al., 2016), we first attempted to develop a pipeline for seed counting that incorporated ImageJ analysis based on segmentation of seed areas. Using our seed scanning setup (see **Methods**), when fewer than 200 seeds were placed on the plate lid and separated using forceps, seeds were detected and counted with high accuracy (correlation between true and predicted seed counts, r^2^, was 0.996, slope=0.9998, 60 images, **Fig. 1b,c, Table S7**). Our ImageJ pipeline allowed the detection of about 52 template images (total of 624 plate lids) per hour with a typical laptop (Intel(R) Core i7-7500U CPU, 16GB RAM).

However, when seeds were placed on plate lids without separation, big clumps of seeds were not counted by ImageJ, and small clumps where a small number of seeds were touching each other were recognized as single seeds (**Fig. 3a**). The prediction accuracy of ImageJ drops off as the number of seeds increases (**Fig. 3c, Table S8**); this is because the more seeds there are on the plate lid, the more likely it is that seeds touch each other, leading to an increase in the false negative rate of prediction. Moreover, the detection of seeds could be disrupted by scratches or letters on the plate lids, and seeds outside the predefined circular search regions were not detected (purple arrowheads in **Fig. S4**). Thus, to obtain accurate counts based on segmentation, it is necessary to separate seeds and confine them to the center of the plate lid, which is time consuming and not amenable to high-throughput analysis.

**Fig. 3.**
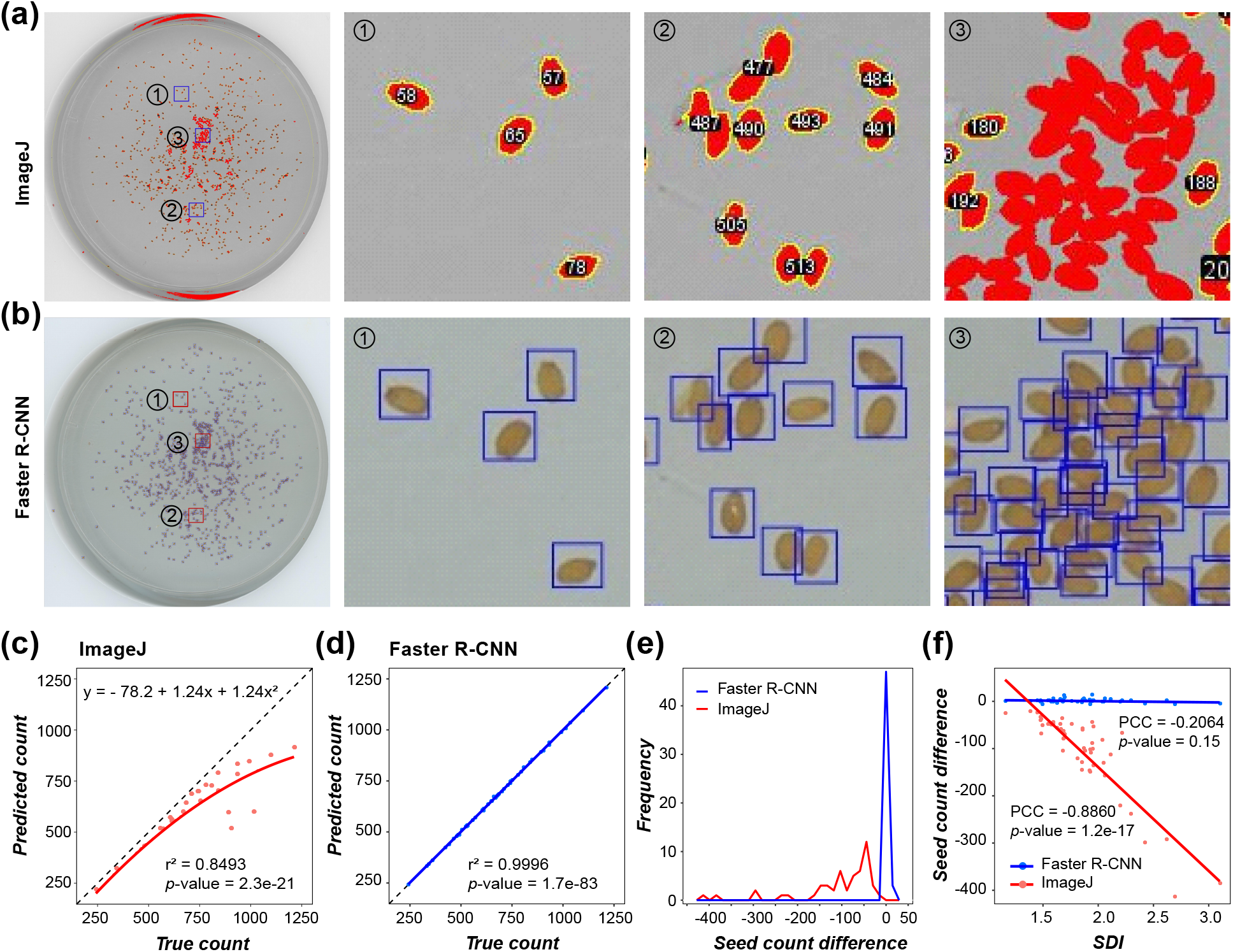
Comparison between the performances of ImageJ- and Faster R-CNN-based seed counting methods for the test set images of seeds that were not deliberately separated. (**a, b**) The same seed scan image analyzed by ImageJ (**a**) and Faster R-CNN (**b**). Three different regions of the plate lid with different densities are outlined. Region 1 has low seed density, region 2 has moderate density, and region 3 has a high density. In (**a**) the red colored regions represent the segmented areas identified by ImageJ; seeds outlined in yellow and assigned numeric IDs were counted. In (**b**) the blue rectangles represent seeds detected by Faster R-CNN. (**c,d**) Correlation between true and predicted seed numbers from ImageJ (**c**) and Faster R-CNN (**d**) analysis of the test set. (**e**) Distribution of differences between true and predicted seed numbers. Red lines: ImageJ; blue lines: Faster R-CNN. (**f**) Correlation between seed density index (SDI) and difference between true and predicted seed counts. Each dot in (**c,d,f**) corresponds to one of the 50 test set images. The red line in (**c**) is the regression line obtained using the loess method. The blue lines in (**d,f**) are fitted regression lines for Faster R-CNN predictions. The red line in (**f**) is the fitted linear regression line for ImageJ-based predictions. PCC: Pearson correlation coefficient.

### Improved seed counting by Faster R-CNN

Next, we evaluated the performance of the deep learning approach Faster R-CNN in seed counting. Since it is very time-consuming to annotate a large number of small objects for model training, we adopted a two-step strategy to reduce the manual labor needed for seed labeling. First, we split the 256 whole-plate images into 1024 quarter-plate images, and manually labeled a subset (180) of these quarter-plate images to speed up the training process. A total of 160 labeled quarter-plate images (*Training image set 1* in **Fig. 2a**) were used to build the models, and the remaining 20 images were set aside as the validation set (*validation image set* in **Fig. 2a**) to evaluate the performance of the models. A model (Model_seed_ 66) built with the optimal hyperparameter combination (scale-B, aspect ratio-A and 10,000 proposals, for details about hyperparameter tuning, see **Methods**) was used to detect seeds in the remaining 844 quarter-plate images to produce “*in silico*” seed annotations for the second-round modeling (**Fig. 2a**). The predicted coordinates of seeds for a set of four quarter-plate images were combined and converted to the corresponding coordinates in the original whole-plate image. These whole-plate seed coordinates were manually corrected (i.e., false negatives were manually labeled, and false positives were removed) to produce a new set of seed annotations, resulting in 211 labeled whole-plate images.

A new model, Model_seed_ 67, with the same parameters as Model_seed_ 66, was built using 161 (*Training image set 2* in **Fig. 2b**) out of these 211 images (**Fig. 2**). The remaining 50 labeled whole-plate images (*Test image set* in **Fig. 2b**) were used to evaluate the performance of Model_seed_ 67, which had an improved average F1 of 0.992 (**Table S8**) compared with the F1 (∼0.970) of Model_seed_ 66 (**Fig. S2**). Note that the test set images were not used for training or validating Model_seed_ 67; they were thus ideal for independently testing the model. In contrast to the ImageJ approach, Model_seed_ 67 correctly predicted seeds regardless of whether they were in contact with each other or not (**Fig. 3b**), and the prediction accuracy was not influenced by the total number of seeds in an image (r^2^=0.9996, *p*=1.7e-83, **Fig. 3d**). The differences between true and predicted seed counts were close to zero, much smaller than those in ImageJ analysis (**Fig. 3e**). Furthermore, Model_seed_ 67 allowed the detection and counting of seeds in about 240 whole-plate images per hour using 1 GPU (Nvidia Tesla K80) with 4 GB of GPU memory in a UNIX cluster, or about 33 images per hour using a laptop with 16 GB of memory. These results suggest that our Faster R-CNN-based models provide highly accurate Arabidopsis seed counts and can be used for large-scale fitness studies.

### Impact of seed density on the Faster R-CNN model

The number of seeds in an image has a detrimental effect on the performance of ImageJ, but not on that of Faster R-CNN, as the correlation between the true and predicted numbers of seeds is nearly perfect (**Fig. 3d**). This suggests that the Faster R-CNN model performance was not affected by the seed density. To verify this, a measure of seed density was needed because the same number of seeds could be evenly distributed on a plate lid, or crowded in a specific area so that seed density was very high in some parts of the lid but low in others. Thus, we generated the seed density index (SDI) as a measure of seed density. First, a circle with a radius of 30 pixels (corresponding to 0.62 mm, approximate length of two seeds) was drawn from the center of a seed, then the number of seeds with central points located within the circle were calculated. Finally, the average number of seeds per circle in a whole-plate image was defined as the SDI (**Fig. 4a**).

**Fig. 4.**
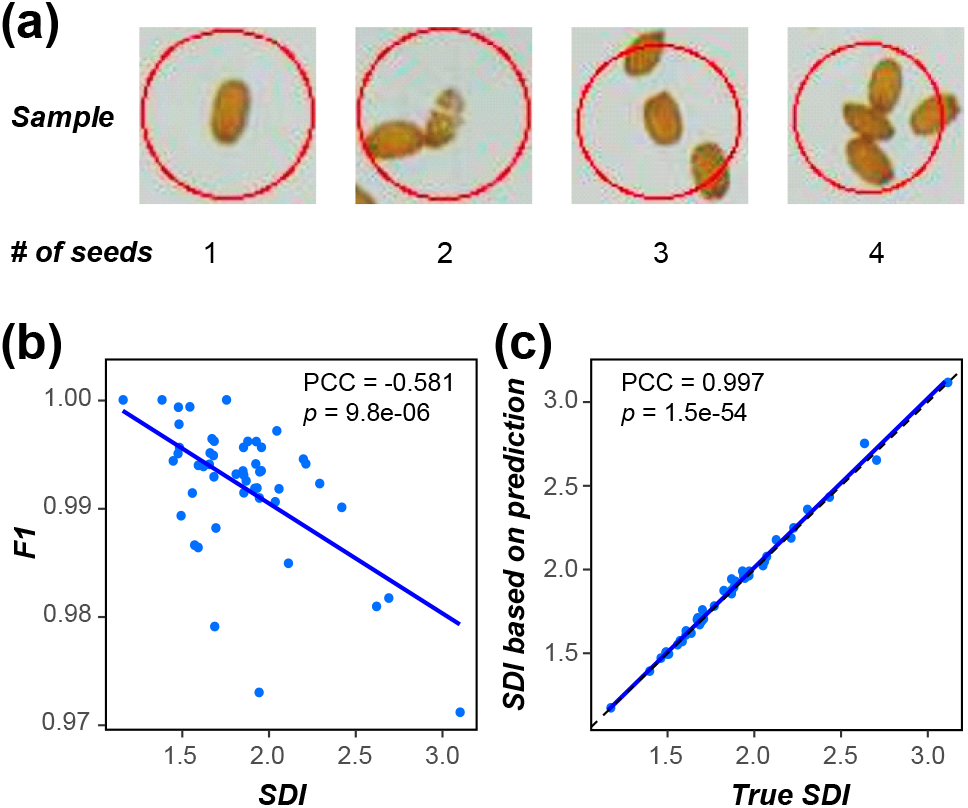
Effect of seed density on the performance of the Faster R-CNN models. (**a**) Examples with different seed density index (SDI) values. The radius of each circle is 30 pixels (0.62 mm). (**b,c**) Relationship between SDI and model performance (**b**) and between the true SDI and SDI based on prediction (**c**) for test images. Each dot corresponds to one of 50 test set images. Blue lines are the fitted linear regression lines. F1: F1 value at 0.5 IoU (Intersection over Union). PCC: Pearson correlation coefficient.

Using this strategy, we calculated the SDIs of the test set images (for examples see **Fig. S5**) and determined the Pearson’s Correlation Coefficient (PCC) between SDI and the performance of Model_seed_ 67 on the test set images (**Fig. 4b**). The higher the seed density, the lower the model performance (PCC between SDI and F1 was −0.581, *p*=9.8e-06, **Fig. 4b**; for the correlation between SDI and other performance measures see **Fig. S6**). Nevertheless, the effect of seed density on the performance of Model_seed_ 67 was small, as the F1 only dropped from 1.000 for an SDI of 1.157 to 0.971 for an SDI of 3.100 (**Fig. 4b, Table S8**). An F1 of 0.971 with a recall of 0.968 indicates that for an image with 1000 seeds, there would only be 32 false negatives (seeds not detected) and 25 false positives (seeds detected in an area with no seeds or a seed area counted more than once). Consistent with this, there was no significant correlation between the SDI and the difference between true and predicted seed counts (PCC=-0.206, *p*=0.15), in contrast to the significant negative correlation observed for ImageJ (PCC=-0.886, *p*=1.2e-17, **Fig. 3f**). We also calculated SDIs for the predicted seed coordinates and found that the PCC value between true and prediction-based SDIs was 0.997 (*p*=1.5e-54; **Fig. 4c**), demonstrating that our Faster R-CNN model also predicts the locations of seeds very well.

### Model improvement through data augmentation

Our goal is to provide a seed counting model that can be widely used by different researchers, who may have seed images with different properties. Thus, we investigated the utility of Model_seed_ 67 for seed counting using images with four varying attributes: resolution, contrast, brightness, and blurriness (**Fig. 5a**). These modified seed images were created by modifying the properties of the test set images (**Fig. 2b,** for the image property settings see **Table S9**). In the modified test set, there were 1750 images: the original test set images (50) and modified images with 34 different attributes (34 × 50, light green box, **Fig. 2b**). A slight but significant decrease in F1 was observed when the brightness of the images was ≤ 0.60 (*p*=0.01, one-sided Wilcoxon signed-rank test) relative to the original images, while the F1 dropped dramatically when the relative brightness was ≥ 1.20 (*p*=6.4e-08, **Fig. 5b**). A significant decrease in F1 was also observed when the relative contrast of images (relative to the original image) was ≤ 0.50 (*p*=1.0e-07) or ≥ 1.75 (*p*=5.0e-4), the relative blurriness was ≥ 1.50 (*p*=6.7e-10), or the relative resolution was ≤ 0.50 (*p*=9.1e-10, **Fig. 5b**). These results suggest that although Model_seed_ 67 is suitable for a range of image qualities, the seed detection accuracy will decrease dramatically when the image properties deviate from the training images beyond a certain point.

**Fig. 5.**
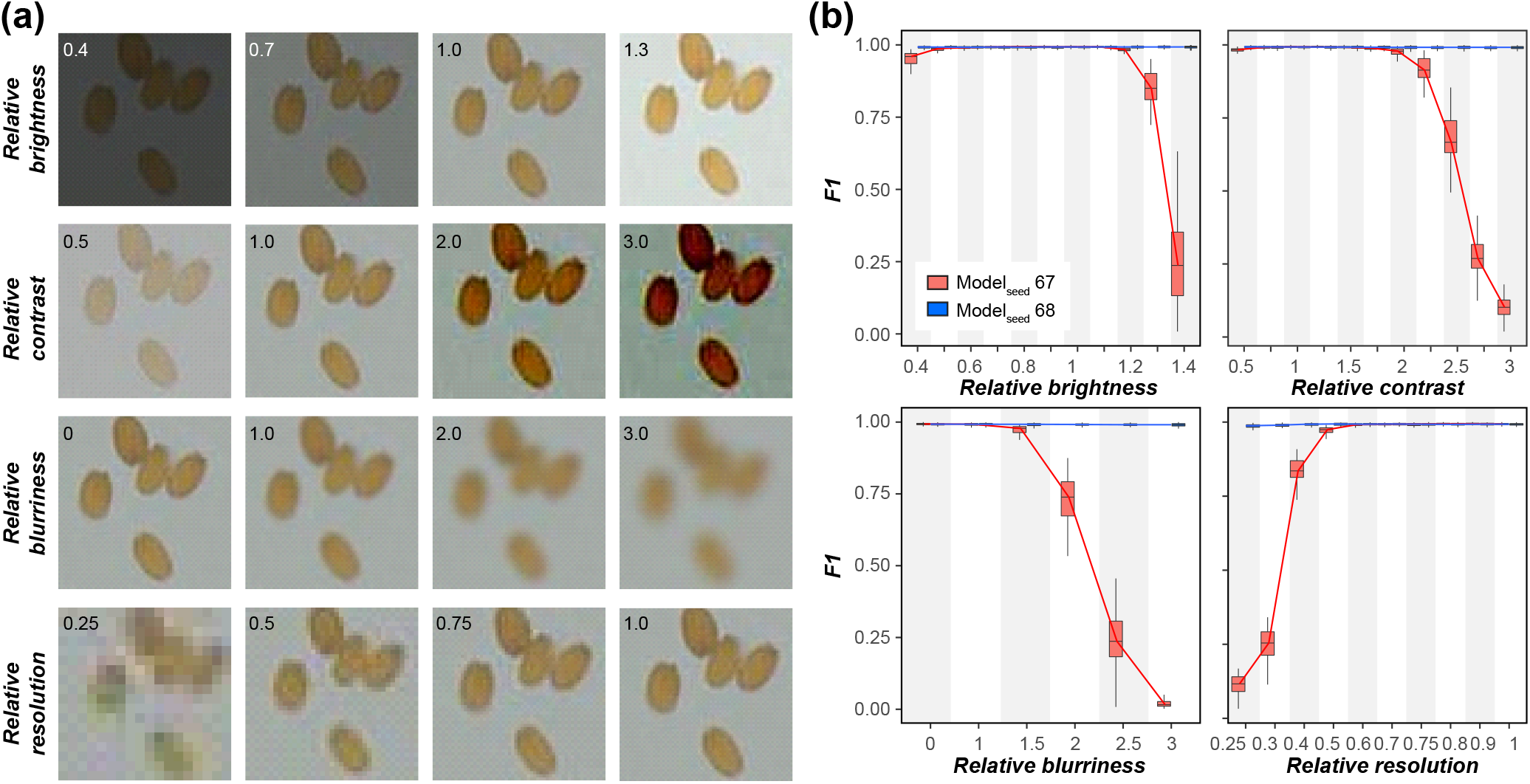
Improvement of model robustness using training images with different properties. (**a**) Examples of seed images with different relative brightness, contrast, blurriness, and resolution values that were derived from the same original image. (**b**) Model performance for Model_seed_ 67 and Model_seed_ 68 on the modified test set (**Fig. 2b**). Red boxplot: Model_seed_ 67; blue boxplot: Model_seed_ 68. F1: F1 value at 0.5 IoU.

To improve the robustness of Model_seed_ 67, we applied data augmentation, in which the size and properties of training datasets are increased so better prediction models can be built (Shorten & Khoshgoftaar, 2019). To accomplish this, we used 20 of the 161 training set 2 images to produce additional images with 21 different property settings (21 × 20, darker green box, **Fig. 3b,** for the image property settings see **Table S9**). These 420 additional images, together with the original 161 images, were used to build a new model, Model_seed_ 68 (**Fig. 2b**), with the same hyperparameter settings as Model_seed_ 67. Model_seed_ 68 was then used to detect seeds in the modified test set images. Although there was a slight decrease in F1 when the relative blurriness was ≥ 3.00 (*p*=0.04, median F1 decrease=0.002) or when the relative resolution was ≤ 0.30 (*p*=0.02, median F1 decrease=0.003, **Fig. 5b**), Model_seed_ 68 (blue, **Fig. 5b**) performed better than the non-augmented Model_seed_ 67 (red, **Fig. 5b**) in all situations and thus, the augmented model is robust to different image properties.

### Fruit counting using Faster R-CNN models

Compared with seed number, total fruit count is an even more frequently used proxy for fitness. Because dry Arabidopsis fruits shatter easily, it is not always possible to harvest all fruits produced by a single plant after seeds have matured, especially for plants growing in the field. In this case, the best method would be to count all fruits (including dehisced ones) and count seeds per fruit for a subset that haven’t dehisced, and then calculate total seed number by multiplying the number of seeds per fruit by the total fruit number. Thus, to obtain more accurate estimates of seed production per plant, it is necessary to record the numbers of both intact and shattered fruits. With these considerations in mind, we developed Faster R-CNN models to count all fruits without harvesting the fruits first. When capturing the images for fruit counting, a pink background was used to maximize the contrast between the background and the dark, dry fruits and the pale replum of shattered fruits that remained after the valves fell from the fruit (**Fig. 6a,d**). Because there were much fewer fruits in each image than there were seeds when performing seed counting, and the fruits were much larger, we manually labeled the fruits in 120 images without using the two-step strategy used for seed counting.

**Fig. 6.**
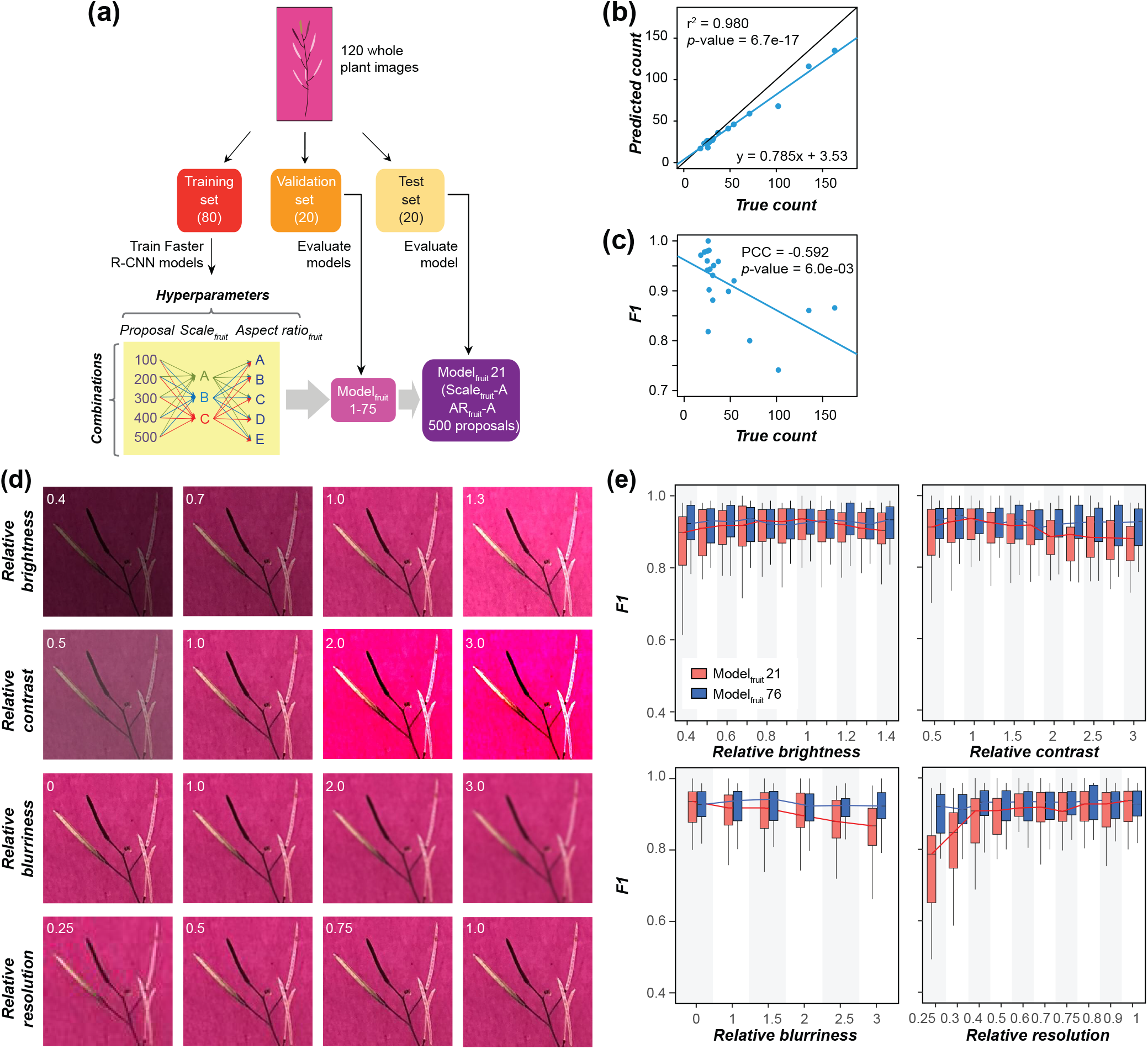
Fruit counting using Faster R-CNN models. **(a**) Fruit counting workflow. (**b**) Relationship between true and predicted fruit numbers. (**c**) Relationship between fruit number in an image and the model performance. PCC: Pearson correlation coefficient. (**d**) Examples of the same fruit image with different relative brightness, contrast, blurriness, and resolution values. (**e**) Model performance for Model_fruit_ 21 and Model_fruit_ 76 on test images with different properties. Red boxplot: Model_fruit_ 21; blue boxplot: Model_fruit_ 76. F1: F1 at 0.5 IoU; PCC: Pearson correlation coefficient.

Eighty, 20, and 20 images were randomly selected and used as training, validation, and test sets, respectively (**Fig. 6a**). Different combinations of hyperparameter values (**Table S6**) were evaluated (Model_fruit_ 1-75, **Fig. 6a**). Surprisingly, all models built with different hyperparameter combinations performed similarly on images in the validation set, with an average F1 of 0.925 (**Fig. S7**). Thus, to minimize the computational cost (lower scales or aspect ratios) while maximizing the number of fruits detected per plant (more proposals), the model built with scale_fruit_-A, aspect ratio_fruit_-A, and 500 proposals (Model_fruit_ 21) was used. Model_fruit_ 21 was applied to the test set images, resulting in an average F1 of 0.914 (**Table S10**). This F1 value translates into 1 false positive and 15 false negatives for an image with 100 fruits. Although the r^2^ between true and predicted fruit counts was 0.980 (*p*=6.7e-17), the detection error increased with an increasing number of fruits in an image and, because the fitted line had a slope of 0.785, the error was mostly due to undercounting or false negatives (**Fig. 6b,c**). Upon further examination, we found that the majority of the false negatives were unopened fruits, especially those that overlapped with the stem or with each other. One potential reason for the failure to detect these fruits is that they are similar to the stem in color and shape. Another reason may be the smaller number of labeled intact fruits (543) compared with the number of pale replums (2082) in our training images. Thus, our model can potentially be improved by increasing the number of intact fruits in the training set.

To assess the robustness of our model on images with different qualities, we applied Model_fruit_ 21 on test set images with different image properties (**Fig. 6d**, modified test set, 700 images, for the image property settings see **Table S9**). Significant decreases in F1 were observed when the relative image brightness was ≤ 0.70 (*p*=0.04) or ≥ 1.40 (*p*=0.02), the relative contrast was ≤ 0.50 (*p*=0.02) or ≥ 1.50 (*p*=0.03), the relative blurriness was ≥ 2.0 (*p*=0.002), or the relative resolution was ≤ 0.6 (*p*=0.05) (**Fig. 6e**). By including images with different properties (**Table S9**) in the training set (1840 images in total), a new model, Model_fruit_ 76, was established and applied to the modified test set. A significant but slight decrease in the resulting F1 values was only observed when the relative resolution was ≤ 0.3 (*p*=0.02, median F1 decrease=0.01) (**Fig. 6e**), indicating the robustness of Model_fruit_ 76. Using this model 180 images could be processed per hour using a UNIX node with 1 GPU and 4 GB graphics memory, and 90 images per hour could be processed using a laptop (1 CPU, 16 GB memory).

### Effects of loss of gene function revealed by measuring fitness traits

To evaluate the importance of fitness traits in investigating gene functions and the utility of our pipeline, the fruits and seeds produced by loss-of-function mutants of three genes, *AtPAP2*, *AtKIN7.4*, and *AtRDK1*, were counted and compared with those of WT. *AtPAP2* was previously reported to modulate carbon metabolism, and overexpression of *AtPAP2* resulted in earlier bolting and a higher seed yield than WT, whereas a loss-of-function mutant of *AtPAP2* (*pap2*) showed no significant differences in plant growth or seed yield relative to WT (Sun et al., 2012). One possible explanation for the failure to detect a difference between *pap2* and WT is genetic redundancy (Sun et al., 2012), but an alternative reason could be that the difference is subtle and not easily detected. To estimate yield, the authors measured seed weight per plant, seed weight per 100 mg of seed, and the number of fruits per plant, and all three of these traits were not significantly different between *pap2* and WT. It is possible that these measurements may not have captured the differences in fitness between *pap2* and WT. To test this, we counted the seeds of *pap2* plants (the same mutant line used by Sun *et al*.) using our seed counting pipeline and observed a significant decrease in seed count compared with WT (*p*=5.1e-4, **Fig. 7a**), consistent with the increased yield phenotype of the overexpression lines described in Sun et al. (2012). This result indicates that the previous failure to detect a decrease in seed yield may have been due to the fitness measure used (i.e., seed weight). In contrast, there was no significant difference in fruit counts between the *pap2* and WT plants (*p*=0.26, **Fig. 7b**).

**Fig. 7.**
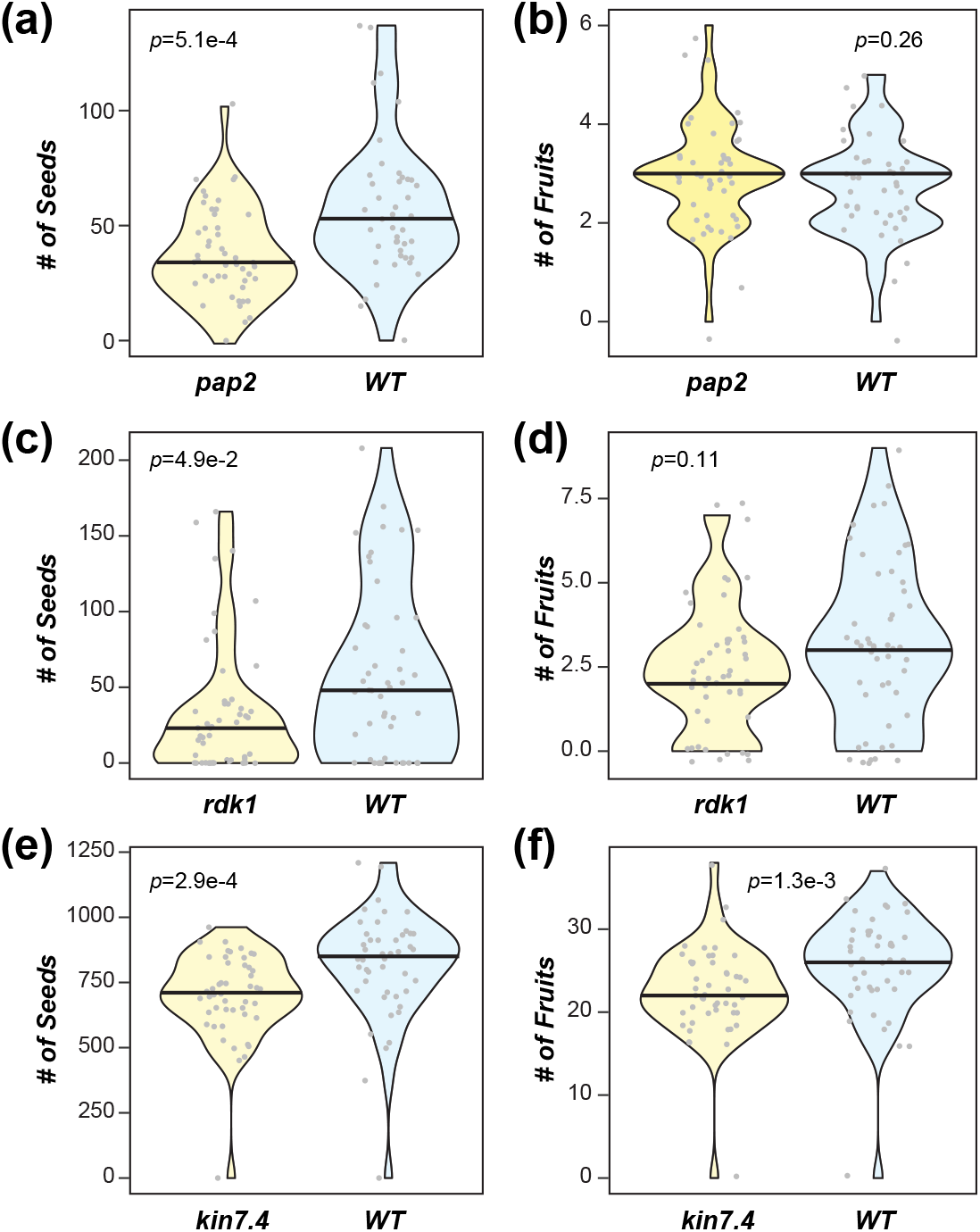
Fitness measurements for three mutants. (**a-b**) Seed (**a**) and fruit counts (**b**) of the T-DNA mutant of *PURPLE ACID PHOSPHATASE 2* (*pap2*; *n* = 47) and wild type (WT; *n* = 43). (**c-d**) Seed (**c**) and fruit counts (**d**) of the T-DNA mutant of *RECEPTOR DEAD KINASE1* (*rdk1*; *n* = 50) and WT (*n* = 50). (**e-f**) Seed (**e**) and fruit counts (**f**) of the T-DNA mutant of *KINESIN 7.4* (*kin7.4*; *n* = 48) and WT (*n* = 46). *p*-values are from Wilcoxon signed-rank tests.

*AtRDK1* is involved in abiotic stress responses and functions in an ABA-dependent manner (Kumar et al., 2017). Phenotypic differences between *rdk1* and WT were previously observed to be condition specific, e.g., differences in cotyledon greening were only observed in the presence of ABA (Kumar et al., 2017). In our study, without imposing any specific treatment, *rdk1* produced significantly fewer seeds than WT (*p*=4.9e-2, **Fig. 7c**), although there was no significant difference in fruit number (*p*=0.11, **Fig. 7d**). This result suggests that fitness may reveal loss-of-function effects that are too subtle to detect or are only detectable under specific conditions.

The third gene we evaluated, *AtKIN7.4*, belongs to the kinesin motor family, members of which are involved in microtubule-based movement (Moschou et al., 2016). No functions have been reported for *AtKIN7.4*. We found that the *kin7.4* mutant produced significantly fewer seeds (*p*=2.9e-4, **Fig. 7e**) and fruits (*p*=1.3e-3, **Fig. 7f**) than WT, indicating that loss of *AtKIN7.4* affects fitness, and in this case, the decrease in fitness can be captured by measuring both fruit number and seed number.

## Discussion

Fitness is one of the best measures of gene functionality because it reflects the ability of a plant to survive and reproduce given all the phenotypic effects of the mutation over the lifetime of the individual. For self-pollinating species such as Arabidopsis, fitness is better assessed by counting the numbers of seeds than fruits, as they more directly reflect the number of offspring and reproductive success. Due to the lack of an effective tool enabling high throughput counting of small seeds en masse, seed counts are often estimated indirectly, for example by dividing the total seed weight per plant by the estimated individual seed weight in the same batch (Cvetkovic *et al*., 2017), or by multiplying the fruit count by the average fruit length (Kerwin *et al*., 2015; Taylor *et al*., 2019). However, these approaches may not yield accurate estimates of seed production due to potential measurement errors and the imperfect correlation between seed number and fruit length (Roux *et al*., 2004). Here, we established a model employing a deep learning approach, Faster R-CNN, to count Arabidopsis seeds—one of the smallest objects analyzed using machine vision to date—with a near perfect accuracy (F1=0.992) using images with multiple different properties or qualities. Our model outperforms another CNN-based approach developed by Toda *et al*. (2020) (F1 of about 0.95), where the detected objects were much larger than Arabidopsis seeds. In particular, the Faster R-CNN-based predictions outperform those of ImageJ, a well-known platform with macros/modules for segmentation and morphology extraction (Schneider *et al*., 2012; Cervantes *et al*., 2016; Vasseur *et al*., 2018). In addition, Faster R-CNN is less time consuming because seeds can be accurately detected without first being separated or confined to predefined regions.

One of the challenges when using deep learning approaches is the requirement for a large number of labeled data (in our case, labeled seeds). To overcome this, we adopted a two-step modeling strategy to reduce the labor needed for seed annotations. In step 1, we split the images and used a subset of the split images to build a preliminary model (F1<0.975) and applied it to the remaining images. While the predictions were not perfect, this step drastically reduced the manual annotations needed because we only needed to correct mis-predictions to boost our seed labels by ∼5 fold (29,360 labels in the first-round, 138,929 labels in the second-round). Using this much larger set of seed labels, new models were built (step 2) that had improved model performance (F1=0.992), indicating the effectiveness of our strategy.

The Faster R-CNN approach also shows promise in fruit detection and counting (r^2^=0.98, slope=0.79). The performance of our fruit counting model was better than that of another recently published CNN-based approach, DeepPod (r^2^=0.90, slope ∼0.70, Hamidinekoo *et al*., 2020). In that paper, the task (i.e., fruit detection) was first divided into four classification tasks: the detection of the tip, body, and base of the fruits and the detection of the stem. The separately detected parts were then joined together as a whole fruit. As the authors noted, this post-processing step affected the final fruit detection performance. In our study, the fruits were labeled and detected as whole objects, thus avoiding the need for post-processing. In addition, different from Hamidinekoo *et al*. (2020), where most fruits and the stems in the images were fresh and green, fruits in our study were dry and light brown to gray, or were shattered with only the pale replum remaining. Thus, our fruit counting approach is expected to be applicable to a wider range of Arabidopsis fruit developmental stages. This is especially important when plants must be grown to maturity, and seed counts are estimated by multiplying the average number of seeds per intact fruit by the total number of fruits (intact and dehisced).

Nevertheless, our fruit counting models did not perform as well as our seed counting models and a published ImageJ-based segmentation and skeletonization approach (r^2^=0.91, slope= ∼1; Vasseur *et al*., 2018), which may be due to the much fewer labeled fruits than labeled seeds (about 52 times more). Thus, the performance of the fruit counting model is expected to be improved when more fruit labels are included to train the model. In addition, one notable drawback of our approach is the undercounting at higher fruit numbers; this was mainly due to overlap between intact fruits and between intact fruits and stems. To remedy this, one approach is to rearrange the inflorescences before capturing the images to keep fruits from overlapping with each other and with stems. Another potential approach is to analyze multiple images (or frames of a movie) taken at different angles or to examine the 3D reconstruction of the inflorescence.

By examining the fitness traits, especially seed counts, we were able to observe phenotypic changes in loss-of-function mutants that were previously not detectable (*pap2*, Sun et al., 2012) or condition specific (*rdk1*, Kumar et al., 2017). In addition, we demonstrated the importance of *AtKin7.4*—the function of which has not been reported previously— for fitness, in terms of both seed and fruit counts. Taken together, our results illustrate the importance of fitness traits in the study of gene functions, and show that Faster R-CNN-based models, which can almost perfectly detect and count Arabidopsis seeds and also detect fruits with high accuracy, are valuable tools in large-scale studies of plant fitness.

## Supplementary Figure Legends

**Fig. S1.** The architecture of Faster R-CNN. The seed images are first processed using a feature extractor (Inception V2), which extracts features from the input images by assigning importance (weights or biases) to the objects in the images. The output of the feature extractor is a feature map, indicating the locations and strength (indicated by color gradient) of the detected features in an image. Then a large number of anchors (rectangles with different aspect ratios [width/height] and scales [relative size]) are generated and placed uniformly throughout the feature map. By applying one of the key modules of Faster R-CNN, Regional Proposal Network, each anchor is assigned an objectness score, which is an indication of how likely it is that the anchor contains an object. A predefined number of anchors (object proposals) are selected based on the rank of objectness scores. Next, to determine whether an anchor contains an object and to adjust the anchors to better fit the location of the seed, the feature maps and the proposals are processed using another module, the Fast R-CNN Detector. Two scores are obtained: a classification score (the likelihood that the proposal region contains a seed) and regression score (the location of the detected seed).

**Fig. S2.** Hyperparameter tuning for seed counting models. (**a-c**) Performance of 63 models (Model_seed_ 1–63) trained on training set 1 with 100 (**a**), 500 (**b**), and 1,000 (**c**) proposals at three scales (columns) and with seven aspect ratios (colored lines; for scale and aspect ratio values see **Table S4**). Performance was evaluated using the validation set. (**d**) Model performance for different proposals based on the scale-B and aspect ratio-A combination. x axis: the number of training steps; y axis: F1 at 0.5 IoU.

**Fig. S3.** Computational efficiency of seed counting models. Computational efficiency of seed counting models trained with training set 1, with 100 (**a**), 500 (**b**), and 1,000 proposals (**c**) at different scales (A, B, and C columns) and with different aspect ratios (colored lines). x axis: the number of training steps; y axis: global steps per second.

**Fig. S4.** Example false negatives from ImageJ analysis and Faster R-CNN models. (**a,b**) One example seed scan image analyzed by ImageJ (**a**) and the Faster R-CNN Model 67 (**b**). Purple arrowheads: example false negatives from ImageJ and Faster R-CNN; green arrowheads: seeds correctly detected by Faster R-CNN.

**Fig. S5.** Example images with different SDI values. Six images of seeds in petri dish lids with SDI values ranging from low (1.157) to high (3.100).

**Fig. S6.** Effect of seed density on the performance of the Faster R-CNN models using different measures of performance. (**a-c**) Relationship between SDI and precision (**a**), recall (**b**), and accuracy (**c**). Each blue dot represents one of the 50 test set images, and lines are the fitted linear regression lines.

**Fig. S7.** Hyperparameter tuning for fruit counting models. Performance of fruit counting models trained on images in the training set with different proposal numbers (rows) at different scales (columns) and with different aspect ratios (colored lines). Performance was evaluated using images in the validation set. For scale and aspect ratio values see **Table S6.** x axis: the number of training steps; y axis: F1 at 0.5 IoU.

## Acknowledgments

We thank Christina B. Azodi, Bethany M. Moore, Siobhan Cusack, and Liang Xu for help in manual seed annotation, and Ally Schumacher for providing the photo of the template. We thank Dirk Colbry for helpful discussions. This work was supported by the U.S. Department of Energy Great Lakes Bioenergy Research Center (BER DE-SC0018409) and the National Science Foundation (DGE-1828149 to SHS; IOS-2107215 to MDL and SHS; DEB-1655630 to PJK, and DEB-1655386 to JKC and SHS).

## Author contributions

PW, FM, JKC, PJK, MDL, and SHS conceived and designed the study. PW, FM, PD, SH, NLP, EV, EW, JKC, PJK, and MDL performed data collection and analysis. PW, FM, MDL, and SHS wrote the manuscript. All authors read and approved the final manuscript.

## Data availability

All the scripts used in this study and the final seed and fruit counting models are available on Github at: https://github.com/ShiuLab/Manuscript_Code/tree/master/2021_Arabidopsis_seed_and_fruit_count

## Supporting information

**Table S1.** Lines used for training seed counting models.

**Table S2.** Lines used for training fruit counting models.

**Table S3.** Lines used for analysis of fitness.

**Table S4.** Hyperparameter space for seed counting.

**Table S5.** Seed counts in 20 quarter-plate images in the validation set.

**Table S6.** Hyperparameter space for fruit (silique) counting.

**Table S7.** Seed counting using ImageJ.

**Table S8.** Seed counting for 50 test set seed images using Model_seed_ 67.

**Table S9.** Image property setting for Model_seed_ 68 and Model_fruit_ 76.

**Table S10.** Fruit counting for 20 test fruit images using Model_fruit_ 21.

