## Supplementary Figures for "High throughput measurement of *Arabidopsis thaliana* fitness traits using deep learning"

### *New Phytologist* Supporting Information

Article acceptance date: Click here to enter a date.

The following Supporting Information is available for this article:

**Fig. S1** The architecture of Faster R-CNN

**Fig. S2** Hyperparameter tuning for seed counting models

**Fig. S3** Computational efficiency of seed counting models

**Fig. S4** Example false negatives from ImageJ analysis and Faster R-CNN models

**Fig. S5** Example images with different SDI values

**Fig. S6** Effect of seed density on the performance of the Faster R-CNN models using different measures of performance

**Fig. S7** Hyperparameter tuning for fruit counting models

**Table S1.** Lines used for training seed counting models.

**Table S2.** Lines used for training fruit counting models.

**Table S3.** Lines used for analysis of fitness.

**Table S4.** Hyperparameter space for seed counting.

**Table S5.** Seed counts in 20 quarter-plate images in the validation set.

**Table S6.** Hyperparameter space for fruit (silique) counting.

**Table S7**. Seed counting using ImageJ.

**Table S8.** Seed counting for 50 test set seed images using Model_seed_ 67.

**Table S9.** Image property setting for Model_seed_ 68 and Model_fruit_ 76.

**Table S10.** Fruit counting for 20 test fruit images using Model_fruit_ 21.

**Fig. S1** The architecture of Faster R-CNN. The seed images are first processed using a feature extractor (Inception V2), which extracts features from the input images by assigning importance (weights or biases) to the objects in the images. The output of the feature extractor is a feature map, indicating the locations and strength (indicated by color gradient) of the detected features in an image. Then a large number of anchors (rectangles with different aspect ratios [width/height] and scales [relative size]) are generated and placed uniformly throughout the feature map. By applying one of the key modules of Faster R-CNN, Regional Proposal Network, each anchor is assigned an objectness score, which is an indication of how likely it is that the anchor contains an object. A predefined number of anchors (object proposals) are selected based on the rank of objectness scores. Next, to determine whether an anchor contains an object and to adjust the anchors to better fit the location of the seed, the feature maps and the proposals are processed using another module, the Fast R-CNN Detector. Two scores are obtained: a classification score (the likelihood that the proposal region contains a seed) and regression score (the location of the detected seed)**.**

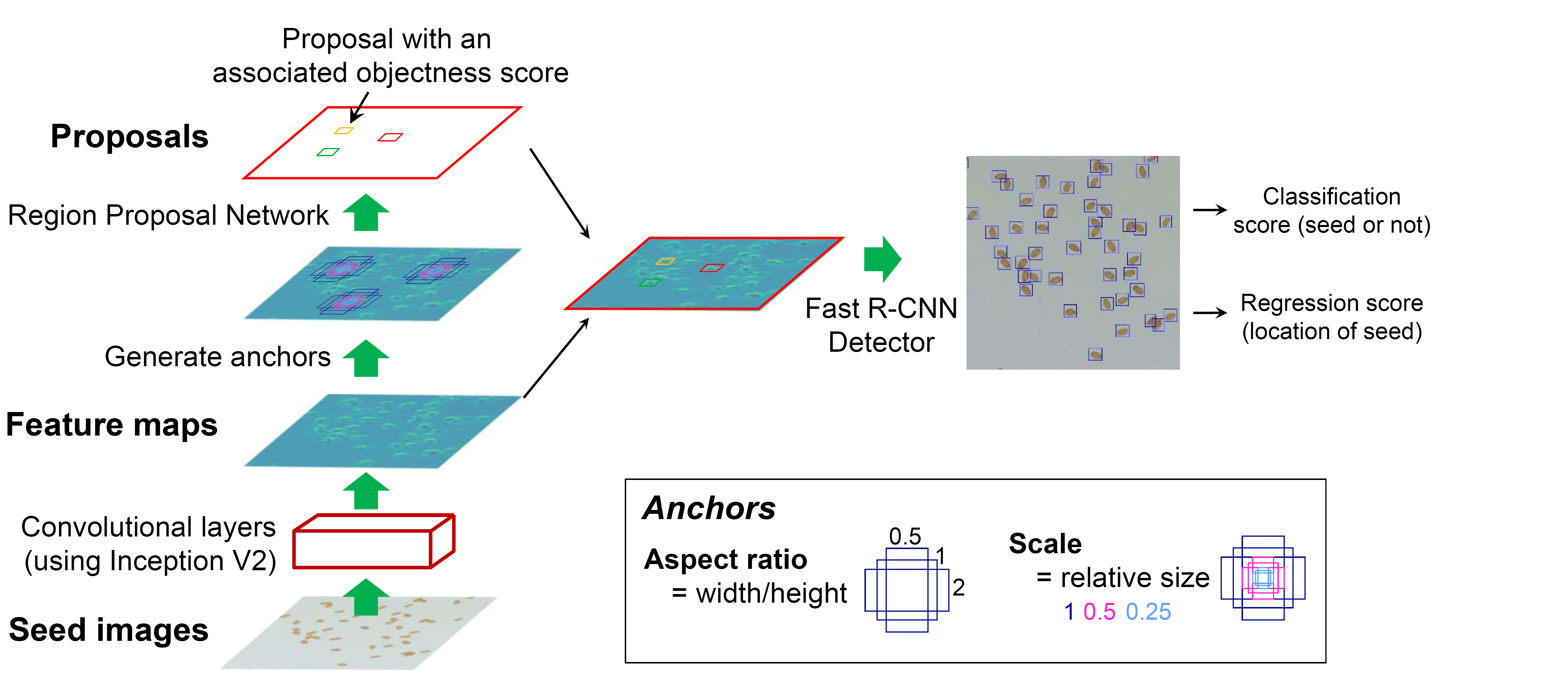

**Fig. S2** Hyperparameter tuning for seed counting models. (**a-c**) Performance of 63 models (Model_seed_ 1–63) trained on training set 1 with 100 (**a**), 500 (**b**), and 1,000 (**c**) proposals at three scales (columns) and with seven aspect ratios (colored lines, for scale and aspect ratio values see **Table S1**). Performance was evaluated using the validation set. (**d**) Model performance for different proposals based on the scale-B and aspect ratio-A combination. x axis: the number of training steps; y axis: F1 at 0.5 IoU.

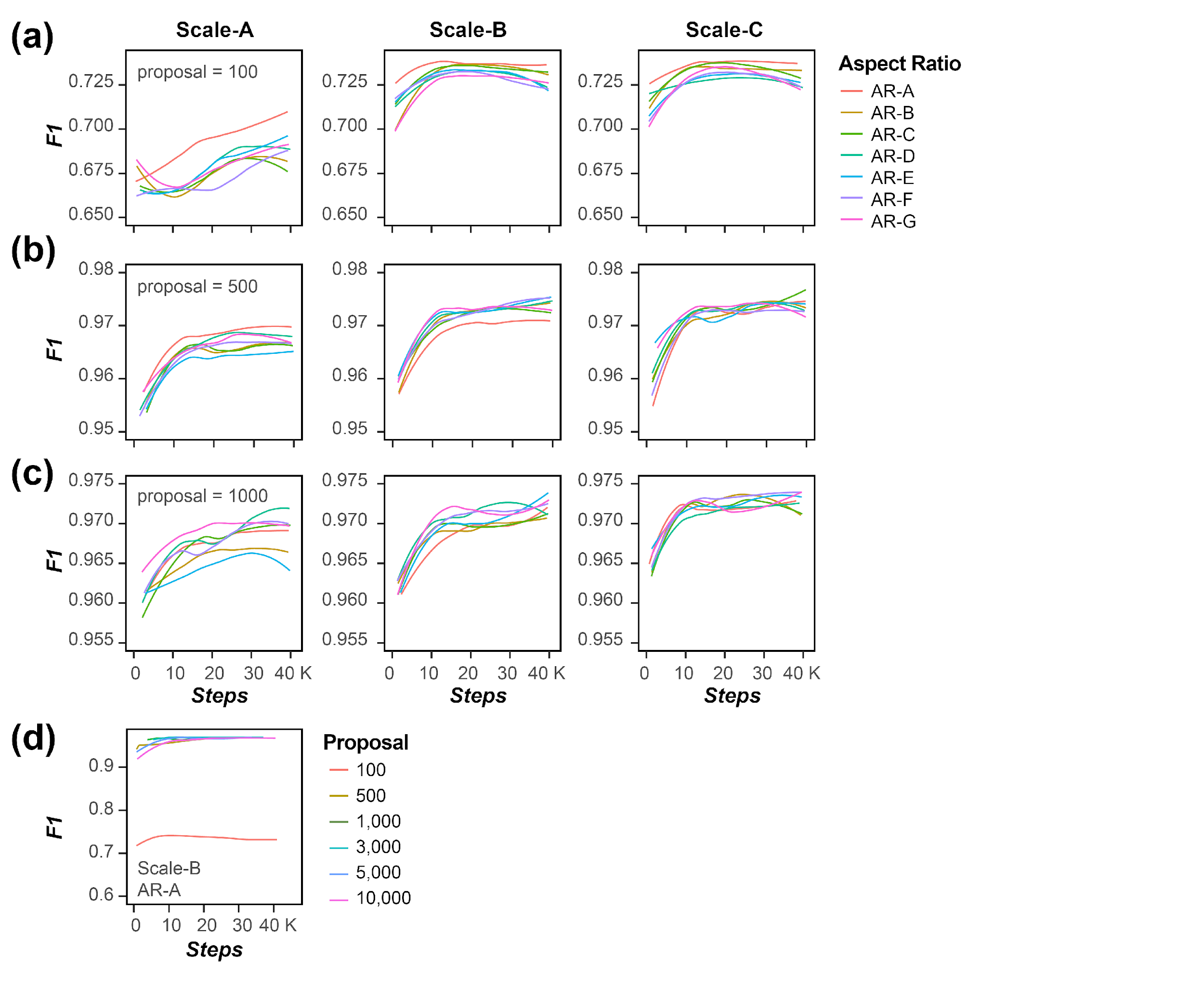

**Fig. S3** Computational efficiency of seed counting models. Computational efficiency of seed counting models trained with training set 1, with 100 (**a**), 500 (**b**), and 1,000 proposals (**c**) at different scales (A, B, and C columns) and with different aspect ratios (colored lines). x axis: the number of training steps; y axis: global steps per second.

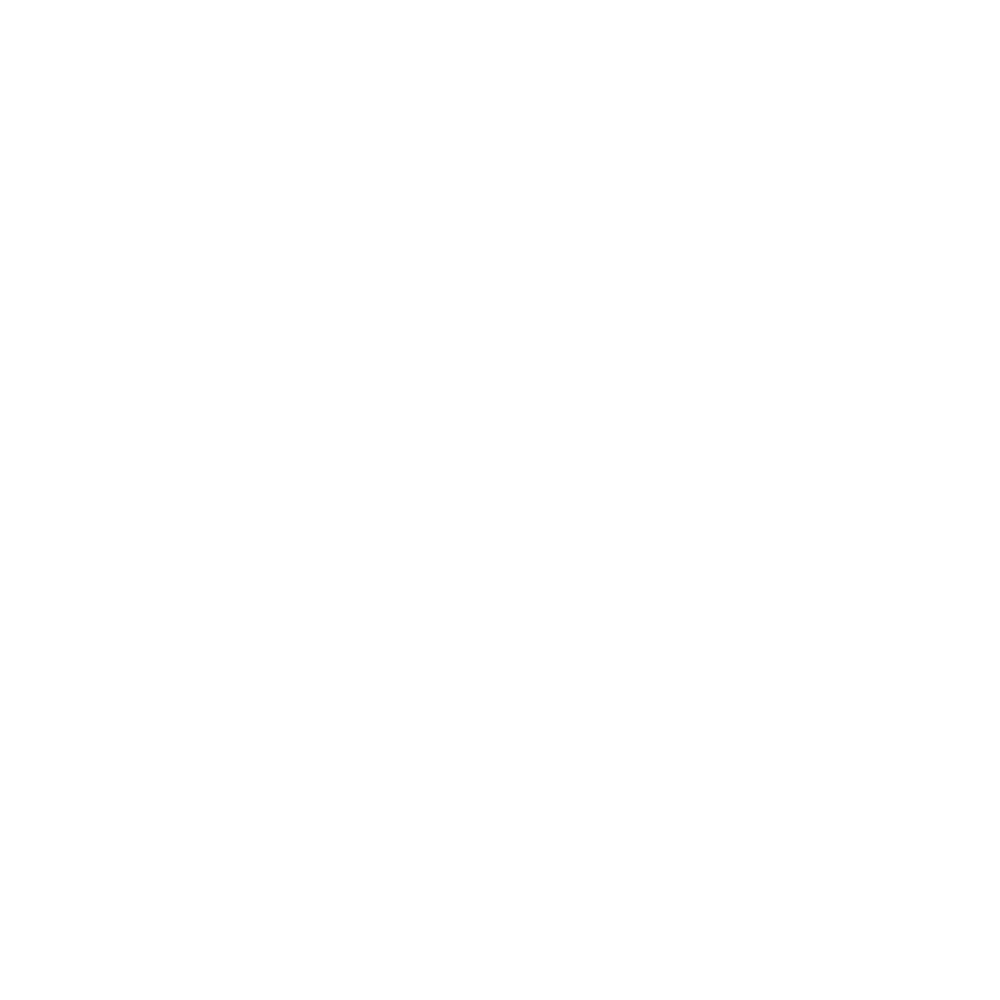

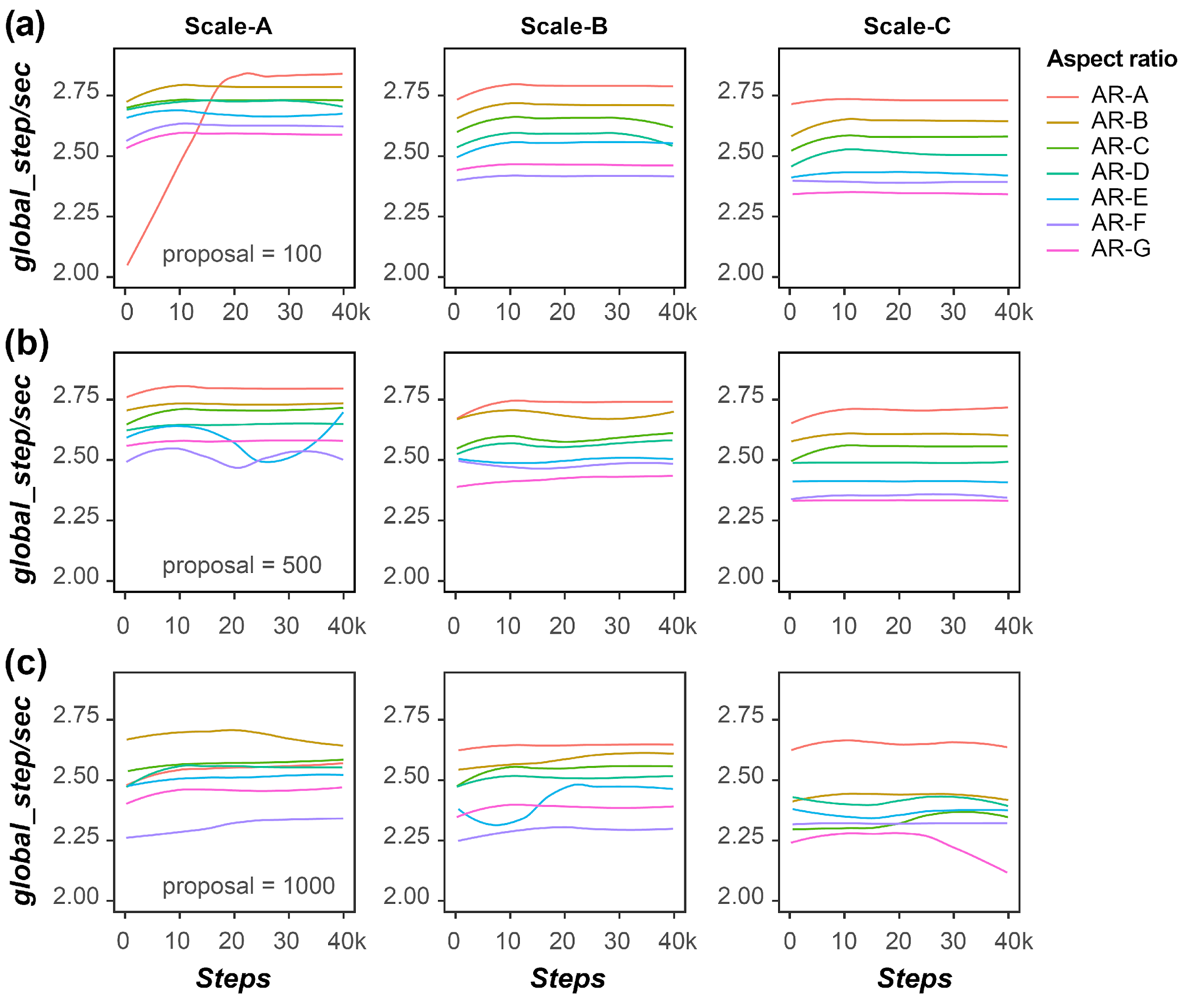

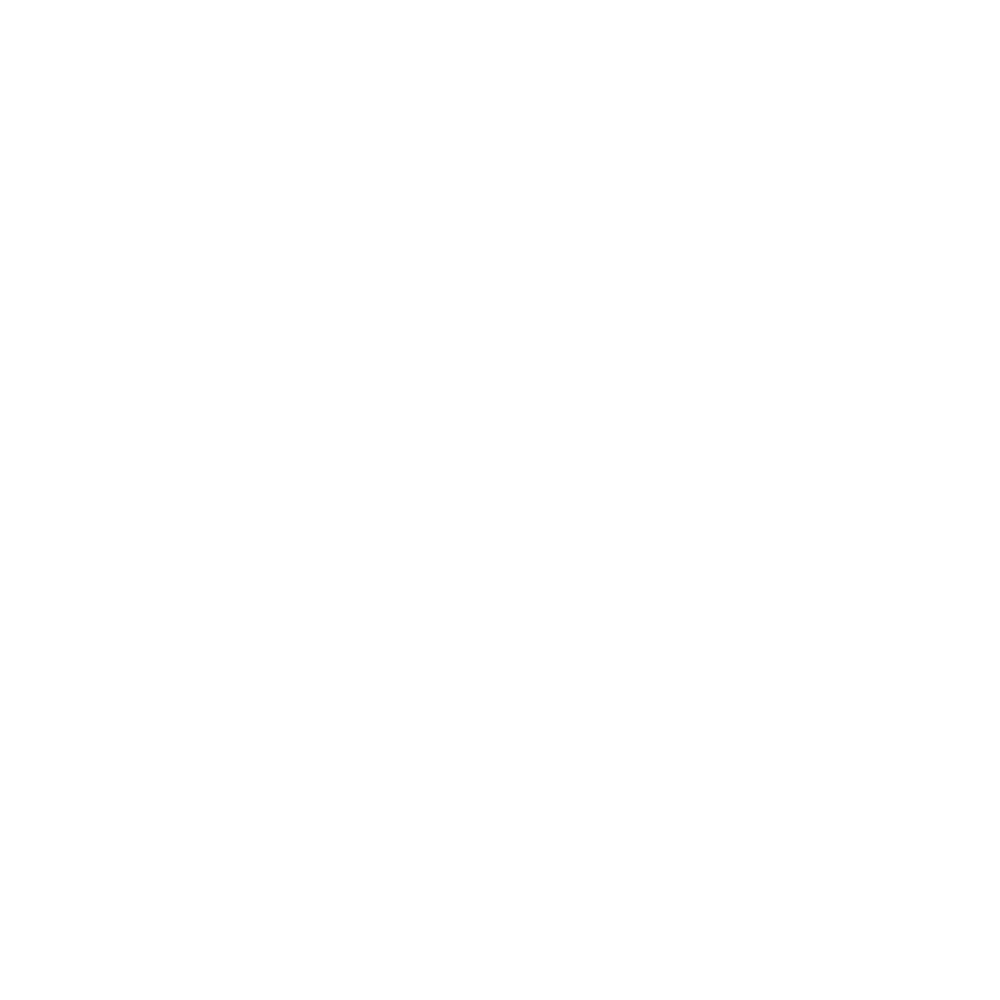

**Fig. S4** Example false negatives from ImageJ analysis and Faster R-CNN models. (**a,b**) One example seed scan image analyzed by ImageJ (**a**) and the Faster R-CNN Model 67 (**b**). Purple arrowheads: example false negatives from ImageJ and Faster R-CNN; green arrowheads: seeds correctly detected by Faster R-CNN.

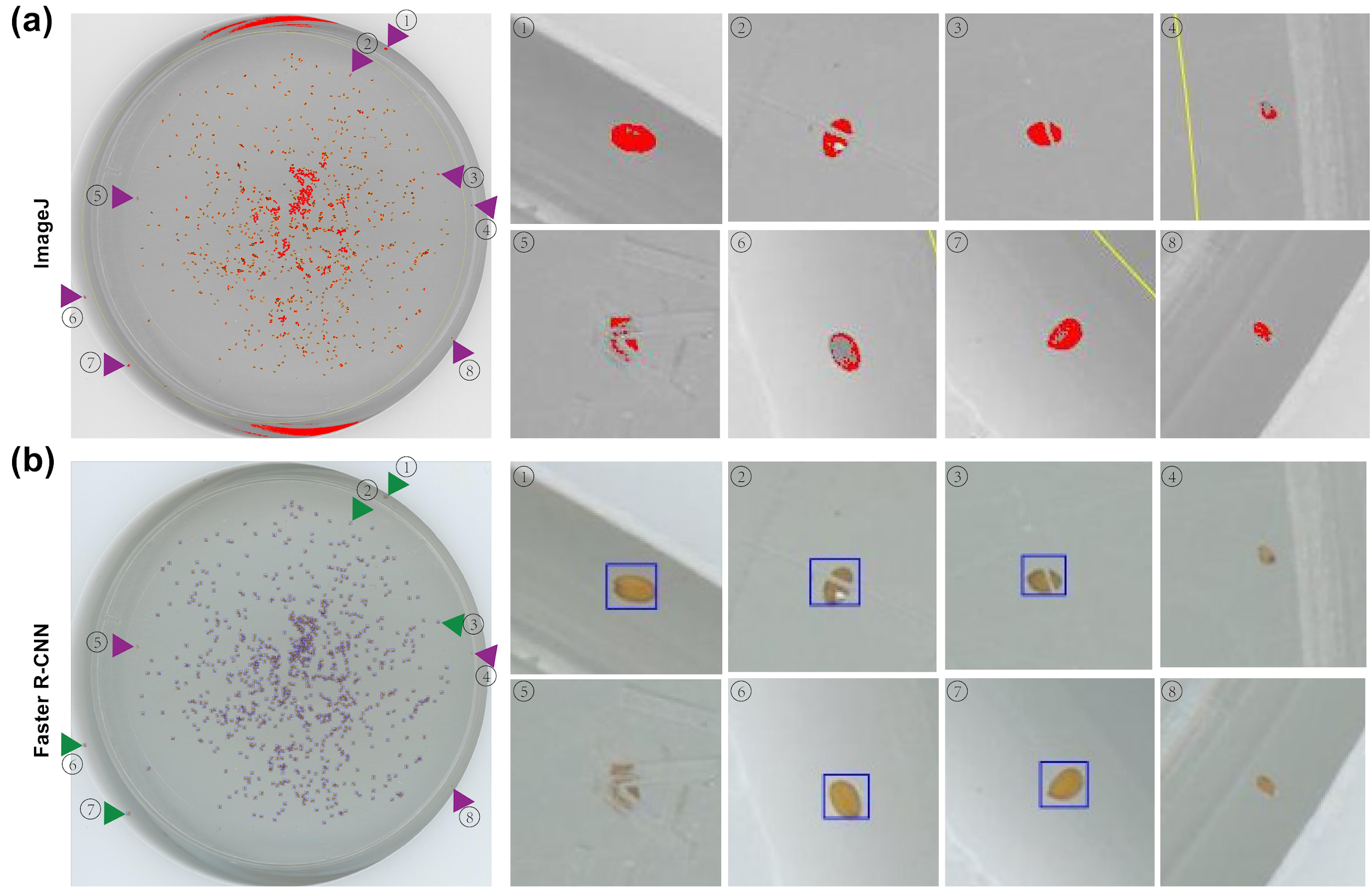

**Fig. S5** Example images with different SDI values. Six seed images with SDI values ranging from low (1.157) to high (3.100).

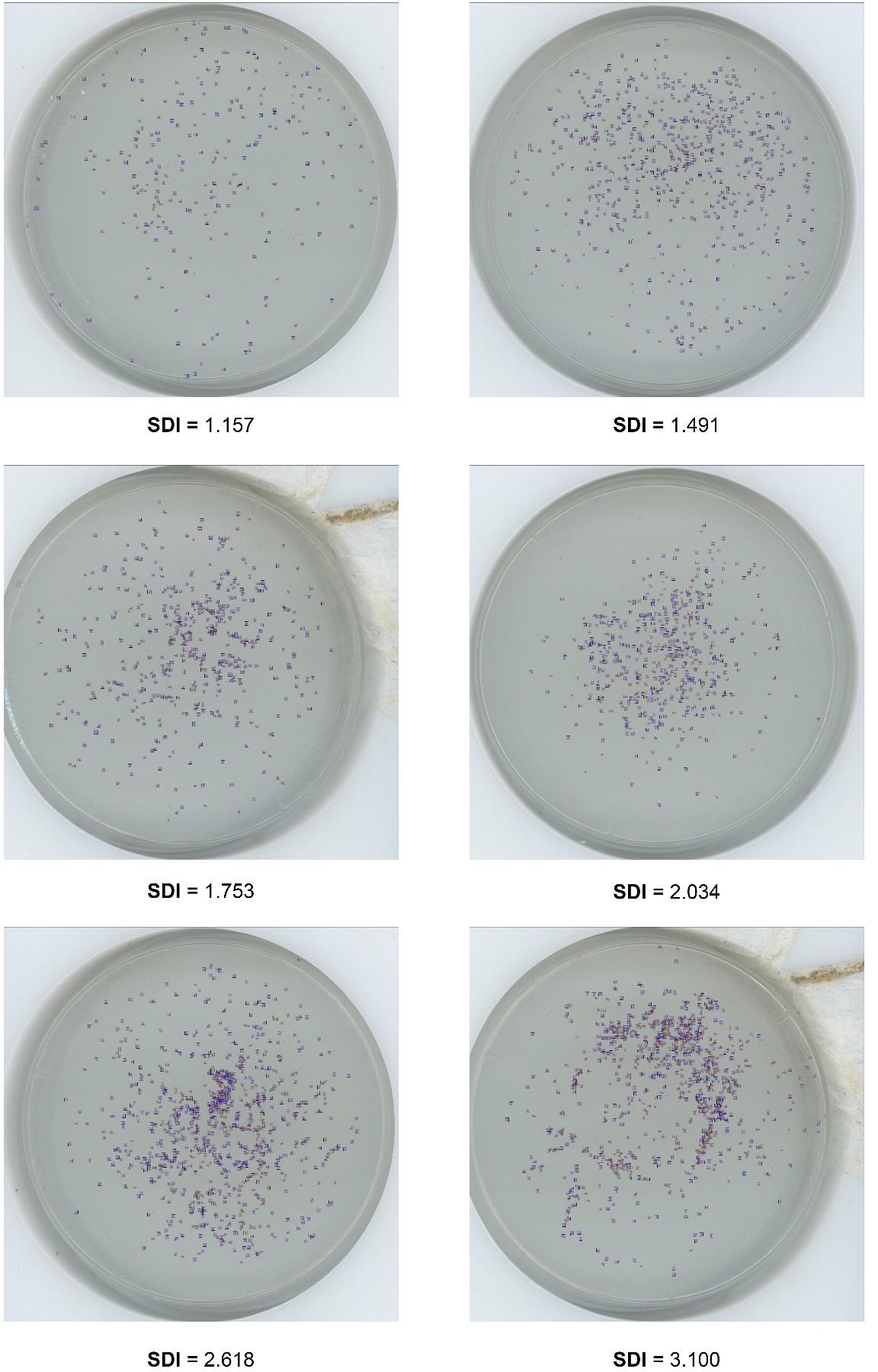

**Fig. S6** Effect of seed density on the performance of the Faster R-CNN models using different measures of performance. (**a-c**) Relationship between SDI and precision (**a**), recall (**b**), and accuracy (**c**). Each blue dot represents one of the 50 test set images, and lines are the fitted linear regression lines.

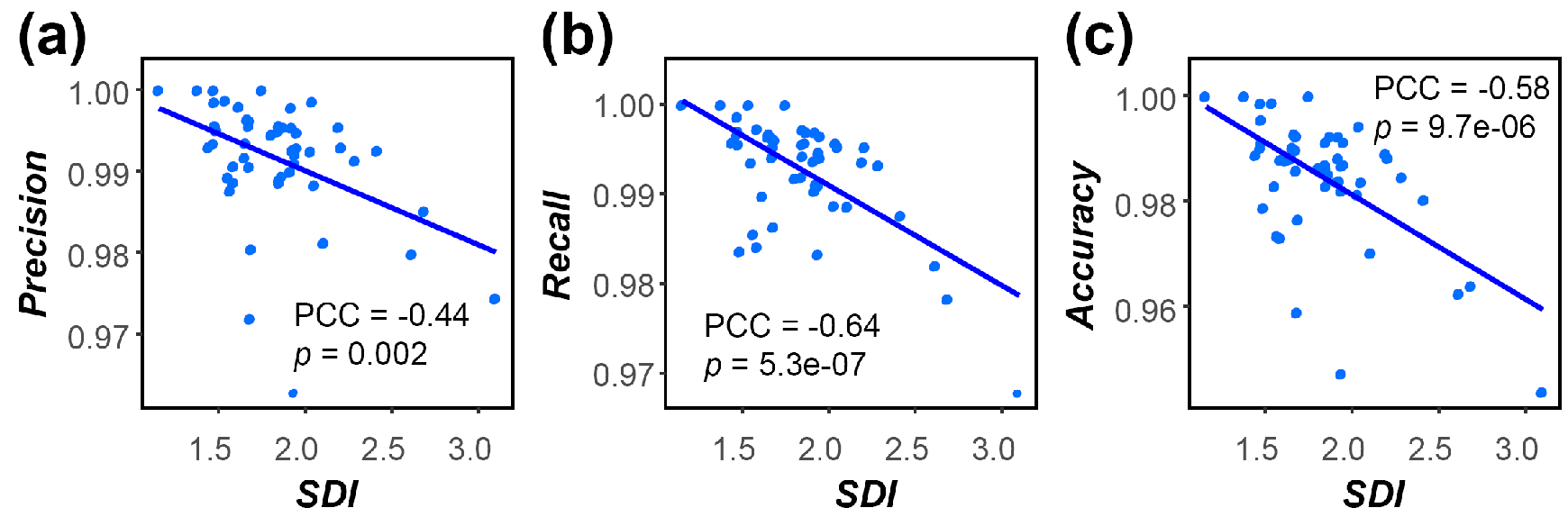

**Fig. S7** Hyperparameter tuning for fruit counting models. Performance of fruit counting models trained on images in the training set with different proposal numbers (rows) at different scales (columns) and with different aspect ratios (colored lines). Performance was evaluated using images in the validation set. For scale and aspect ratio values see **Table S2.** x axis: the number of training steps; y axis: F1 at 0.5 IoU.

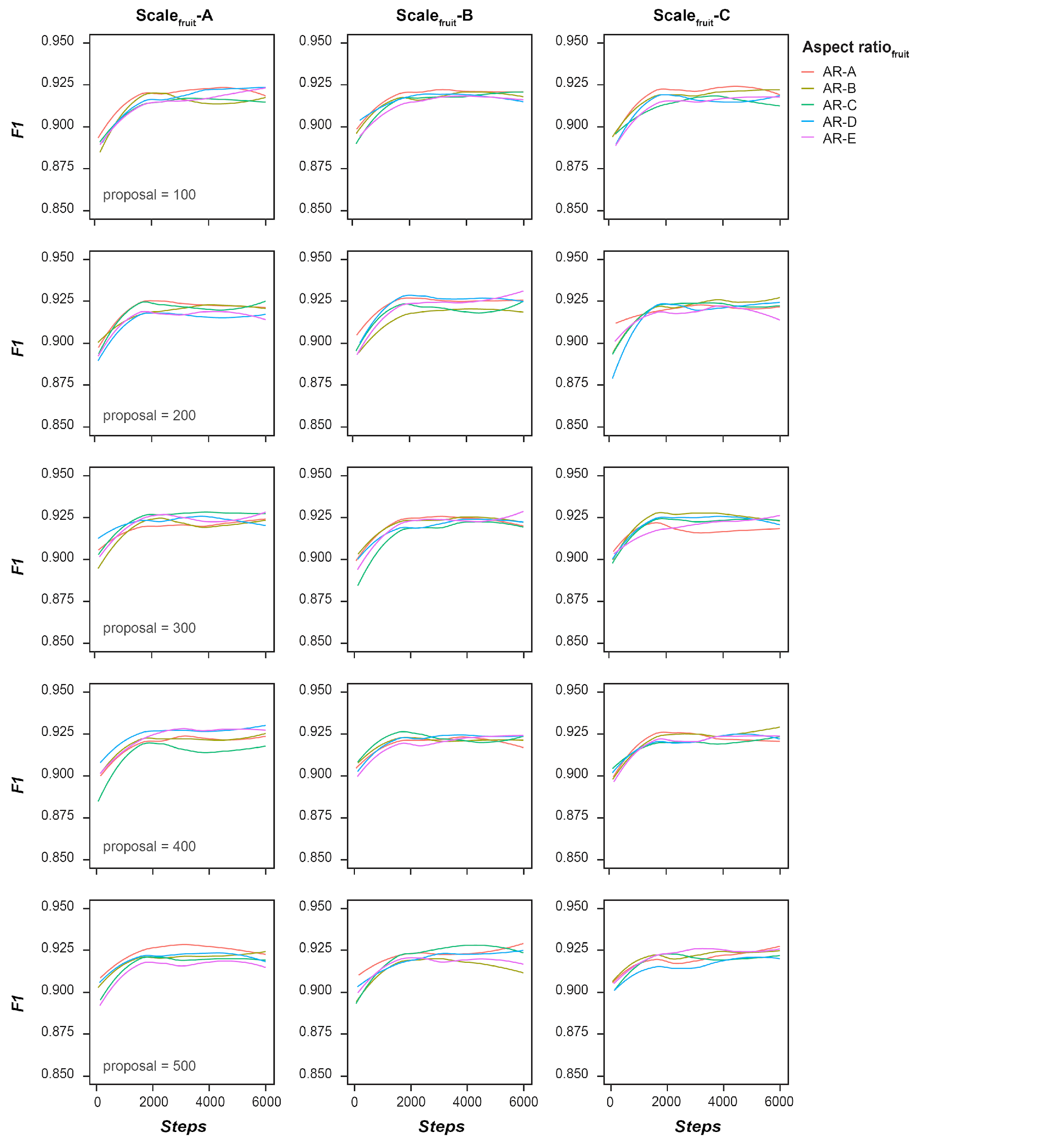
